# DNMT3A clonal hematopoiesis-driver mutations induce cardiac fibrosis by paracrine activation of fibroblasts

**DOI:** 10.1101/2023.01.07.521766

**Authors:** Mariana Shumliakivska, Guillermo Luxán, Inga Hemmerling, Wesley Tyler Abplanalp, Xue Li, Marina Scheller, Carsten Müller-Tidow, Florian Leuschner, Bianca Schuhmacher, Alisa Debes, Simone-Franziska Glaser, Marion Muhly Reinholz, Klara Kirschbaum, Jedrzej Hoffmann, Eike Nagel, Valentina O. Puntmann, David John, Sebastian Cremer, Andreas M. Zeiher, Stefanie Dimmeler

## Abstract

Hematopoietic mutations in epigenetic regulators like DNA methyltransferase 3 alpha (DNMT3A) drive clonal hematopoiesis of indeterminate potential (CHIP) and are associated with adverse prognosis in patients with heart failure (HF). The interactions between CHIP-mutated cells and other cardiac cell types remain unknown.

Here, we identify fibroblasts as potential interaction partners of CHIP-mutated monocytes using combined transcriptomic data from peripheral blood mononuclear cells of HF patients with and without CHIP and the cardiac tissue. We demonstrate that CHIP augments macrophage-to-cardiac fibroblasts interactions. Mechanistically, the secretome of *DNMT3A*-silenced monocytes leads to myofibroblast activation, partially through epidermal growth factor (EGFR) signaling. Harboring DNMT3A CHIP-driver mutations is associated with increased cardiac interstitial fibrosis in mice and patients, and, thereby, may contribute to the poor outcome.

These findings not only identify a novel pathway of DNMT3A CHIP-driver mutation-induced instigation and progression of HF, but may also provide a rationale for the development of new anti-fibrotic strategies.

**Graphical abstract:** 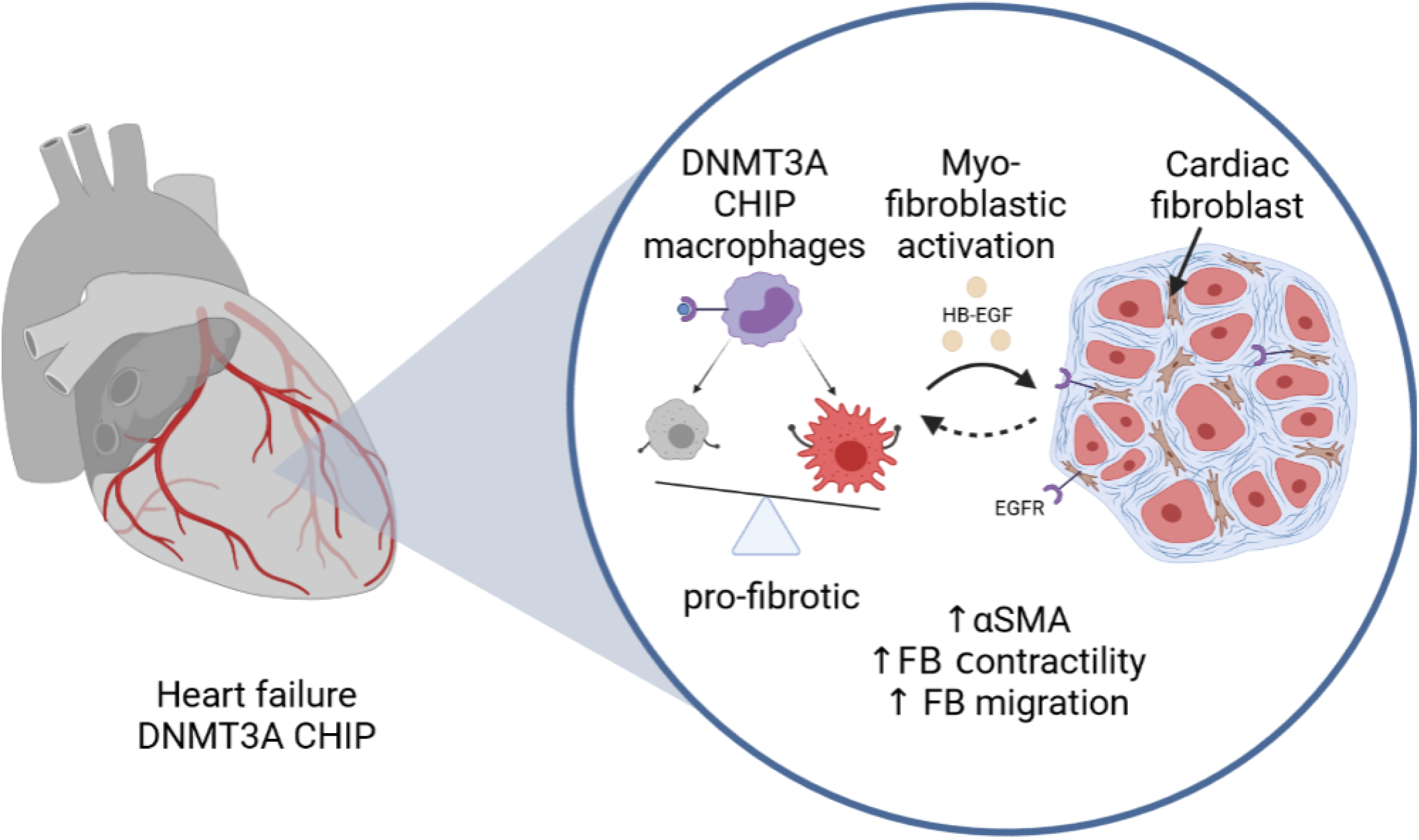

## Introduction

Clonal hematopoiesis of indeterminate potential, or CHIP, is a common age-related condition, in which acquired mutations in hematopoietic stem cells lead to the expansion of a genetically distinct subpopulation of blood cells^1^. The incidence of clonal hematopoiesis increases with age and was previously shown to associate with a variety of cardiovascular diseases, including heart failure^2–7^. Mutations in the epigenetic modifiers DNA methyltransferase 3A (*DNMT3A*) and ten eleven translocation 2 (*TET2*) are the most common CHIP-driver mutations^2^. Importantly, previous studies demonstrated that harboring either DNMT3A or TET2 CHIP-driver mutations in circulating blood cells confers increased mortality in patients with established heart failure as well as aortic valve stenosis^8–12^. Mechanistically, experimental as well as clinical single-cell RNA sequencing studies revealed that DNMT3A and TET2 CHIP-driver mutations regulate the inflammatory potential of circulating immune cells, which may contribute to the promotion of diffuse cardiac fibrosis and, thereby, affect long-term outcome^4,12–16^.

However, as CHIP is defined as the presence of a mutation with a variant allele frequency of at least 2% resulting in at least 4% of circulating myeloid cells harboring the mutation^1^, it still remains a matter of discussion, how such a minor fraction of circulating blood cells may activate inflammatory processes in cardiac tissue leading to subsequent promotion of diffuse cardiac fibrosis and affecting prognosis in patients harboring these mutations. We hypothesized that the interaction between circulating mutant cells, which are continuously recruited to the heart, and cardiac fibroblasts may play a prominent role to aggravate diffuse cardiac fibrosis, thereby promoting disease progression and increased mortality in patients with established heart failure.

## Results

### Interaction of DNMT3A CHIP monocytes with cardiac fibroblasts

To determine a putative interaction of monocytes with cardiac cell types, we integrated single-cell RNA sequencing data of circulating monocytes derived from patients with chronic heart failure with reduced ejection fraction (HFrEF) carrying DNMT3A CHIP-driver mutations (N=5) or no-CHIP carriers (N=4)^15^ with publicly available single nuclei RNA sequencing data of human heart tissue (N=14)^17^ (Fig. 1a). Unsupervised clustering after combining monocytes from PBMC and human heart tissue data sets revealed 9 distinct clusters (Fig. 1b, Supplementary Fig. 1a, b).

**Fig. 1:**
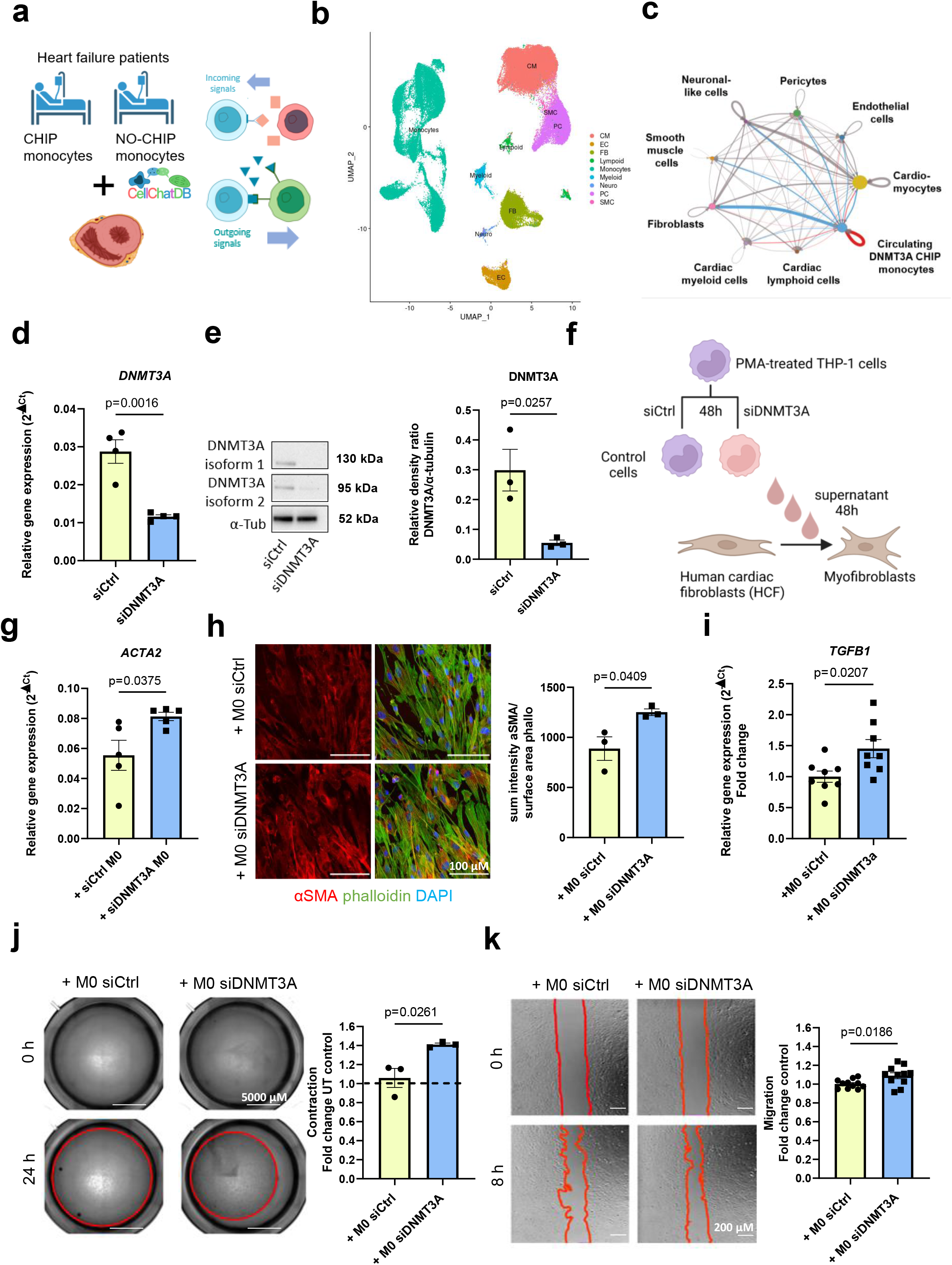
DNMT3A CHIP monocytes activate cardiac fibroblast in a paracrine manner. **a** Schematic representation of the design of a bioinformatic analysis using monocytes derived from scRNA-seq dataset of PBMC from CHIP and No-CHIP patients^15^ and snRNA-seq dataset of cardiac tissue from the septum of 14 control hearts^17^. **b** Representative uniform manifold approximation and projection (UMAP) plot after snRNA-seq and scRNA-seq datasets integration. A total of 9 clusters were identified and annotated as shown in Supplementary Fig.1a-b **c** Circular plot visualizing ligand-receptor interactions determined by DB CellChat of peripheral blood monocytes from CHIP patients with cardiac cell types. Highlighted in blue are predicted interactions between monocytes of CHIP patients and the respective cell types of the cardiac tissue. The thickness of the line indicates the strength of the interaction. **d** Relative *DNMT3A* transcript expression in PMA-activated THP-1 cells after siRNA silencing of *DNMT3A* relative to negative control (n=4) **e** Relative DNMT3A protein expression in PMA-activated THP-1 cells after siRNA silencing of DNMT3A relative to negative control (n=3) **f** Schematic representation of the indirect co-culture experiment. **g** Relative *ACTA2* mRNA expression in human cardiac fibroblasts (HCF) after incubation with supernatants from PMA-activated THP-1 cells for 48 hours after siRNA silencing of *DNMT3A* relative or negative control (n=5). **h** Left, representative immunofluorescence images of αSMA protein expression in stimulated HCF. Blue, DAPI; green, phalloidin; red, αSMA. Right, quantification of αSMA expression (n=3). **i** Relative *TGFB1* mRNA expression in HCF after incubation with supernatants of THP-1 cells (n=8). **j** Left, representative brightfield images of collagen gel contraction in stimulated HCF. Right, quantification of migration (n=3). **k** Left, representative brightfield images of migration in stimulated HCF. Right, quantification of migration normalized to the control (n=11). Data are shown as mean ± SEM. Normal distribution was tested with the Shapiro-Wilk test or the Kolmogorov-Smirnov test. Statistical comparison of two groups was performed using unpaired, two-sided Student’s t-test.

Pooled analysis of cellular interactions by CellChat^18^ demonstrated that monocytes derived from patients carrying DNMT3A CHIP-driver mutations showed highest cumulative interaction strength with cardiac fibroblasts in terms of paracrine signaling compared to all other cell clusters, as indicated by the width of the blue line (Fig. 1c, Supplementary Fig. 1c). These results predict that monocytes, after being recruited to the heart, may interact predominantly with cardiac fibroblasts.

To test this hypothesis, we silenced DNMT3A in PMA-activated THP1 monocytes by siRNAs, which efficiently reduced DNMT3A mRNA and protein expression compared to control siRNAs (Fig. 1d, e). Silencing of DNMT3A with siRNA mimics the most common loss-of-function CHIP mutations^19^. Then, we added the supernatants of the transfected monocytes to cardiac fibroblasts and assessed the effect on fibroblast gene expression and myofibroblast activation (Fig. 1f). Indeed, after treatment with supernatants of DNMT3A-silenced monocytes, fibroblasts showed increased expression of an α-smooth muscle actin (*ACTA2*), which is a common sign of myofibroblast activation^20,21^, on mRNA and protein level (Fig. 1g, h). Moreover, supernatants of siRNA DNMT3A-treated monocytes increased TGFβ expression in fibroblasts (Fig. 1i), which is the prototypical fibrogenic factor in the heart ^22,23^. In addition, we then determined effects of the treatment on fibroblast functions. Supernatants of DNMT3A-silenced monocytes stimulated contraction and migration of cardiac fibroblasts (Fig. 1j, k), which are known features of myofibroblast activation^24,25^. Proliferative capacity of treated fibroblasts showed a trend to increase (Supplementary Fig. 1d).

To further gain insights in how monocytes derived from patients harboring DNMT3A CHIP-driver mutations affect fibroblasts and other cardiac cells in a more physiological environment, we used 3D humanized cardiac tissue mimetics (cardiospheres)^26^ (Fig. 2a). Functionally, supernatants of DNMT3A-silenced monocytes reduced the frequency of contraction of cardiospheres (Fig. 2b-c), while the size of cardiospheres remained unchanged (Fig. 2b, d), suggesting that DNMT3A silencing has no direct effect on cardiomyocyte hypertrophy in this setting. Moreover, PDGFRα-positive area was significantly increased (Fig. 2e), whereas vascularization was not affected (Supplementary Fig. 2a). Consistent with the results of the 2D culture, supernatants of DNMT3A-silenced monocytes also increased α-smooth muscle actin expression (αSMA) (Fig. 2f). Taken together, these data demonstrate that the reduction of DNMT3A in monocytes can induce cardiac fibroblast activation and reduce contraction frequency of cardiac tissue mimetics in a paracrine manner.

**Fig. 2:**
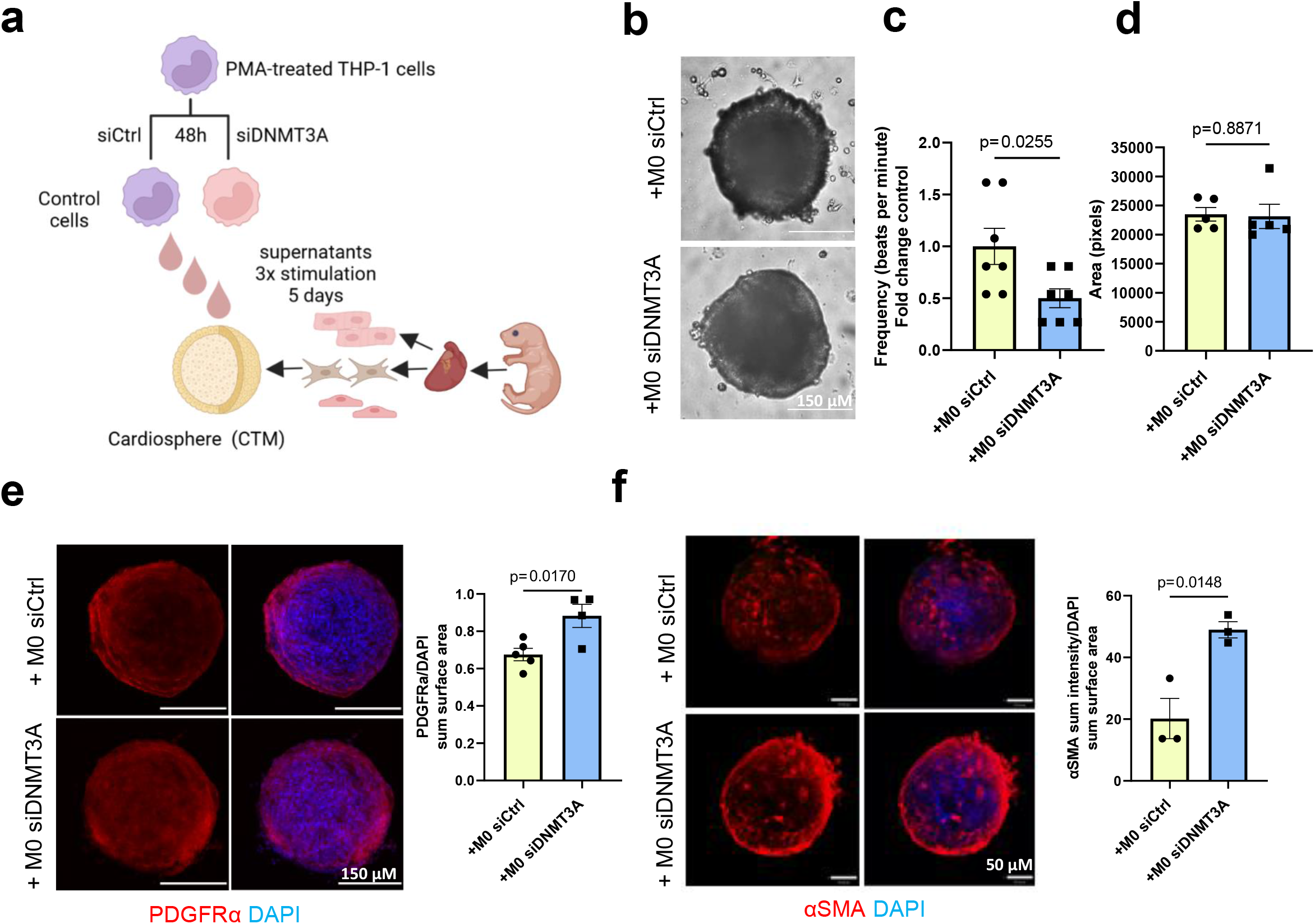
DNMT3A CHIP monocytes reduce contractility and lead to fibrosis in cardiospheres. **a** Schematic representation of experimenal design testing the paracrine effects of THP-1 cells on cardiac tissue mimetics (CTMs, cardiospheres) with THP-1 derived supernatants after siRNA silencing of DNMT3A. **b** Representative brightfield images of cardiospheres upon indirect co-culture (5 days, 3 stimulations). **c** Estimation of beating frequency of cardiospheres stimulated with THP-1 derived supernatants after *DNMT3A* silencing (n=7). **d** Cardiospheres area upon stimulation with THP-1 derived supernatants after *DNMT3A* silencing (n=5). **e** Left, representative immunofluorescence images of PDGFRa protein expression in stimulated cardiospheres. Blue, DAPI; red, PDGFRα. Right, quantification of PDGFRα expression (n=4). f. Left, representative immunofluorescence images of αSMA protein expression in stimulated cardiospheres. Blue, DAPI; red, αSMA. Right, quantification of αSMA expression (n=3). Data are shown as mean ± SEM. Normal distribution was tested with the Shapiro-Wilk test or the Kolmogorov-Smirnov test. Statistical comparison of two groups was performed using unpaired, two-sided Student’s t-test.

### DNMT3A CHIP promotes diffuse cardiac fibrosis in mice and patients with HF

To investigate if DNMT3A CHIP-driver mutations in hematopoietic cells stimulate cardiac fibroblasts *in vivo*, we utilized Dnmt3a^fl-R882H^ mice that express hDNMT3a-R882H in a Cre-dependent manner^27^. Bone marrow cells of pI:pC treated Mx-Cre^+^/Dnmt3a and Mx-Cre^-^/Dnmt3a donor (Ly5.2) mice were transplanted in wildtype recipients (Ly5.1). Myocardial infarction was induced by permanent ligation of the left anterior descending artery^28^ and hearts were analyzed four weeks following surgery (Fig. 3a). Though the scar size in the mice carrying a human DNMT3a-R882H mutation in the bone marrow-derived hematopoietic cells was not significantly affected (Fig. 3b), there was an increase in cardiac interstitial fibrosis in the remote zone of myocardial tissue (Fig. 3c). These data suggest that mimicking DNMT3A CHIP in mice induces diffuse interstitial cardiac fibrosis.

**Fig. 3:**
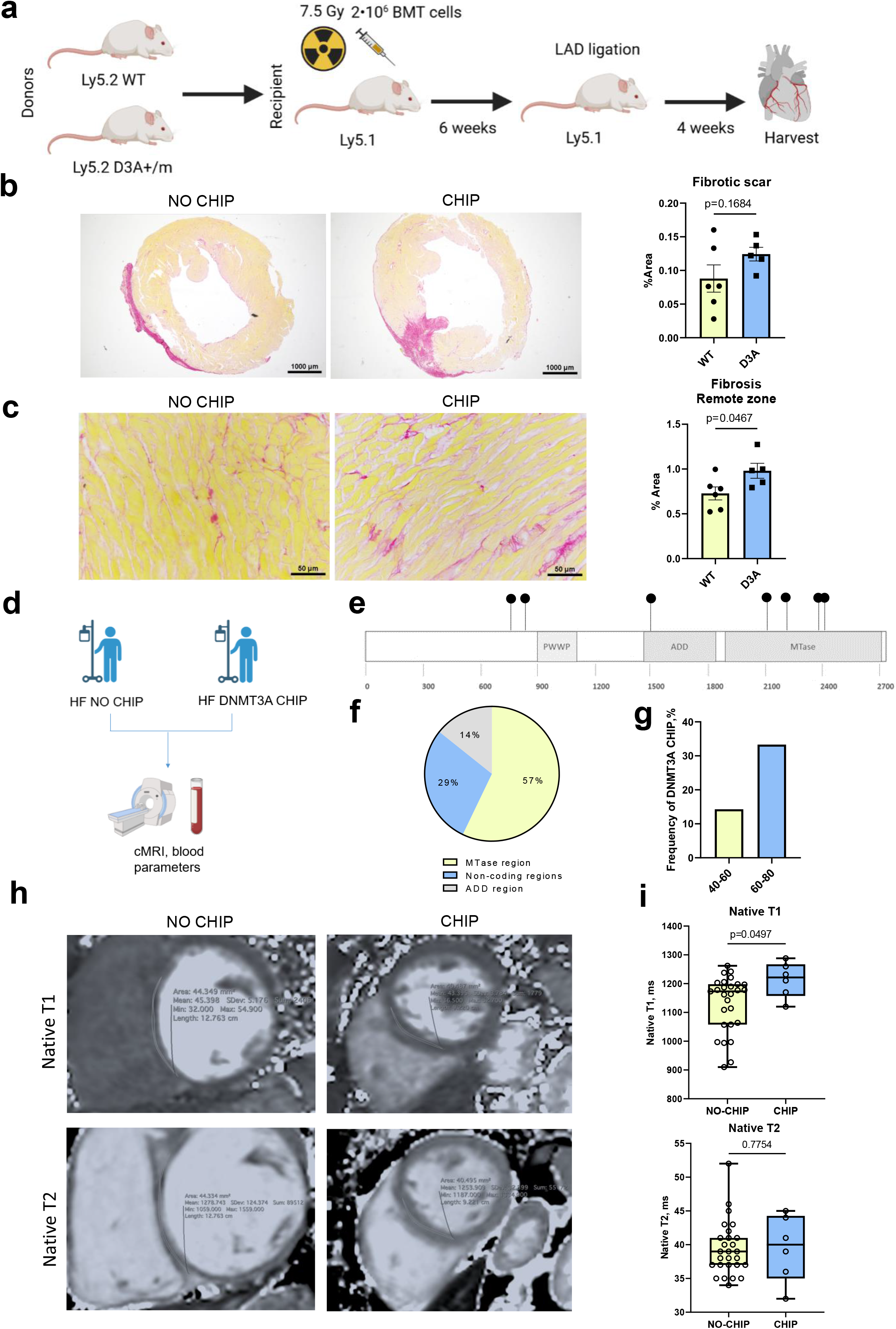
DNMT3A CHIP promotes diffuse cardiac fibrosis in mice and patients with HF. **a** Schematic illustration of the mouse experiment testing the effect of DNMT3A CHIP mutation in hematopoietic cells on cardiac fibrosis. Briefly, bone marrow transplantation of WT (NO CHIP) and D3A+ (CHIP) donor mice into WT mice was performed. After reconstitution of the bone marrow, mice were exposed to LAD ligation and the heart was harvested after 28 days. **b** Left, representative brightfield images of collagen deposition in fibrotic scar in No CHIP and CHIP mice. Right, quantification of fibrotic scar (n=5-6). **c** Left, representative brightfield images of diffuse interstitial cardiac fibrosis in the remote zone in No CHIP and CHIP mice. Right, quantification of the diffuse cardiac fibrosis (n=5-6). **d** Schematic illustration of the design of the clinical study. CHIP and No-CHIP patients were undergoing cardiac magnetic resonance imaging (cMRI) and blood parameters measurement. **e** Distribution of 7 detected mutations of 6 CHIP patients on the scheme of DNMT3A gene; PWWP - Pro-Trp-Trp-Pro motif domains, ADD - ATRX-DNMT3-DNMT3L domain and MTase - methyltransferase domain. **f** Percentage distribution of the CHIP mutations according to the DNMT3A domains. **g** Frequency of DNMT3A CHIP in relation to patient’s age. **h** Representative cMRI sequences of native T1 and T2 relaxation times (ms) in No CHIP and CHIP HF patients. **i** Quantification of native T1 and T2 relaxation times (ms) in No CHIP and CHIP patients where high-quality cMRI sequences were available (n=27 and n=6). Data are shown as mean ± SEM or box plots with min-max values whiskers. Normal distribution was tested with the Shapiro-Wilk test or the Kolmogorov-Smirnov test. Statistical comparison of two normaly distributed groups was performed using unpaired, two-sided Student’s t-test; for the data not following Gaussian distribution Wilcoxon-Mann-Whitney test was applied.

Therefore, we further investigated the impact of harboring DNMT3A CHIP-driver mutations on diffuse cardiac fibrosis using cardiac magnetic resonance (CMR) imaging with myocardial mapping^29–31^ in patients with heart failure (Fig. 3d). 6 patients were harboring DNMT3A CHIP-driver mutations (Supplementary Fig. 2b), but did not differ with respect to sex, age or co-morbidities from No-CHIP patients (Fig. 2e-g, Supplementary Table 1). Strikingly, carriers of DNMT3A CHIP-driver mutations had significantly increased native T1, but no differences in native T2 measurements, indicating the presence of diffuse myocardial fibrosis in these patients (Fig. 3h, i).

### EGF signaling contributes to CHIP monocyte-mediated cardiac fibroblast activation

Next, we assessed the mechanism by which monocytes obtained from DNMT3A CHIP-driver mutation carriers stimulate fibroblast activation. Therefore, we identified the receptor-ligand interactions that were predicted between monocytes derived from DNMT3A CHIP-driver mutation carriers versus NO-CHIP patients and cardiac tissue using CellChat. Epidermal growth factor (EGF) and transforming growth factor β (TGFβ) pathways were among the specifically enriched interactions between monocytes of DNMT3A CHIP-carriers and human cardiac cell types (Fig. 4a). Furthermore, gene ontology term analysis of interactions specific for monocytes obtained from DNMT3A CHIP-driver mutation carriers to cardiac fibroblasts revealed an enrichment of EGF receptor (EGFR) binding and activation of known EGF downstream signaling pathways like MAP kinases activity (Fig. 4b). Since EGFR signaling is a well-known mediator of cardiac fibroblast activation^32,33^, these data suggest that monocytes obtained from DNMT3A CHIP-driver mutation carriers may release factors from the EGF family to induce cardiac myofibroblast activation, which express high levels of *EGFR* in human heart (Fig.4c).

**Fig. 4:**
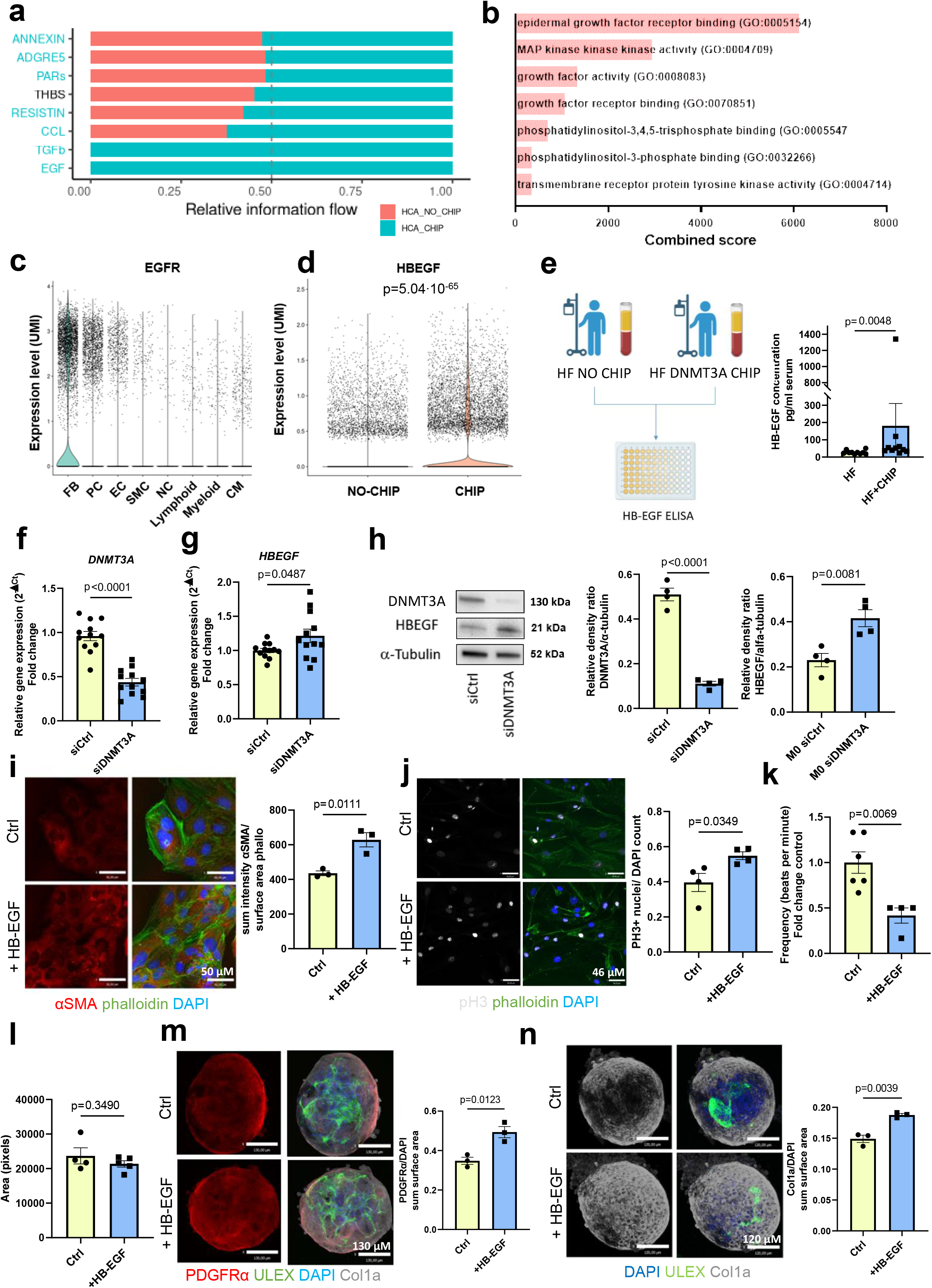
EGF signaling contributes to CHIP monocyte-mediated cardiac fibroblast activation. **a** Signalling pathways dominating relative information flow of CHIP patients monocytes and cardiac tissue based on CellChat cellular crosstalk analysis. **b**. Representation of the most significant functional categories for genes upregulated in CHIP-specific monocytes-to-fibroblasts signaling. Representation was revealed by gene ontology analysis using the Enrichr data base. Graphs represent Enrichr combined score that combines P value and Z score. **c** Violin plots for *EGFR* gene expression in cardiac cell types from snRNA-seq data of cardiac tissue from the septum of 14 control hearts^17^. **d** Violin plot showing *HBEGF* gene expression from scRNA-seq data of monocytes from No-CHIP and CHIP HF patients^15^ **e** Left, illustration of the design of the study. Right, ELISA-based quantification of HB-EGF in serum of the NO-CHIP and CHIP HF patients (n=10 and n=10). **f** Relative *DNMT3A* transcript expression and **g** *HBEGF* transcript expression in PMA-activated THP-1 cells after siRNA silencing of DNMT3A relative to negative control (n=12) **h** Relative DNMT3A and HB-EGF protein expression in PMA-activated THP-1 cells after siRNA silencing of DNMT3A relative to negative control (n=4). **i** Left, representative immunofluorescence images of αSMA protein expression in HB-EGF stimulated iHCF. Blue, DAPI; green, phalloidin; red, αSMA. Right, quantification of αSMA expression (n=3). **j** Left, representative immunofluorescence images of phospho-histone H3 protein expression in HB-EGF stimulated HCF. Blue, DAPI; green, phalloidin; white, pH3. Right, quantification of a pH3 positive nuclei (n=4). **k** Estimation of beating frequency of cardiospheres stimulated with HB-EGF (n=4-6). **l** Cardiospheres area upon stimulation with HB-EGF (n=4-5). **m** Left, representative immunofluorescence images of PDGFRα protein expression in HB-EGF stimulated cardiospheres. Blue, DAPI; red, PDGFRα green, ULEX. Right, quantification of PDGFRα expression (n=4). **n** Left, representative immunofluorescence images of Col1a protein expression in HB-EGF stimulated cardiospheres. Blue, DAPI; white Col1a; green, ULEX. Right, quantification of PDGFRa expression (n=4). Data are shown as mean ± SEM. Normal distribution was tested with the Shapiro-Wilk test or the Kolmogorov-Smirnov test. Statistical comparison of two normaly distributed groups was performed using unpaired, two-sided Student’s t-test; for the data not following Gaussian distribution Wilcoxon-Mann-Whitney test was applied. Violin plots represent log2-transformed and normalised UMI counts. Adjusted p values is based on Bonferroni correction.

CellChat predicted two unique significantly upregulated ligand-receptor pairs between monocytes obtained from DNMT3A CHIP-driver mutation carriers and human cardiac fibroblasts, namely heparin-binding EGF (HB-EGF) with EGFR and amphiregulin (AREG) with EGFR. To test the hypothesis if EGFR signaling is involved in CHIP-driven activation of fibroblasts, we determined the expression of predicted ligands from EGF family members in monocytes obtained from DNMT3A CHIP-driver mutation carriers on single-cell level. The expression of both, *HBEGF* and *AREG*, was significantly higher in circulating monocytes of heart failure patients carrying DNMT3A CHIP-driver mutations compared to NO-CHIP carriers (Fig. 4d, Supplementary Fig. 2c). Moreover, HB-EGF protein was increased in plasma samples obtained from patients with HF harboring DNMT3A CHIP-driver mutations compared to NO-CHIP HF patients as assessed by ELISA (Fig. 4e; Supplementary Table 2). The regulation of HB-EGF expression by DNMT3A was further supported by demonstrating that DNMT3A silencing in monocytes *in vitro* induced overexpression of HB-EGF mRNA and protein levels (Fig. 4f-h). In addition, we observed increased expression of metalloproteases such as *ADAM8* and *ADAM9*, which are known to shed HB-EGF^34^, in DNMT3A CHIP monocytes in patients (Supplementary Fig. 2d) and upon *DNMT3A* silencing *in vitro* (Supplementary Figure 2e). Together these data demonstrate that DNMT3A inactivation leads to an increased expression and possibly activation of HB-EGF in cell culture and in humans.

To gain insights into the functional consequences of the increased HB-EGF expression, we tested the effects of recombinant HB-EGF on cardiac fibroblasts and cardiospheres. Recombinant HB-EGF increased αSMA in cardiac fibroblasts (Fig. 4i), increased proliferation (Fig. 4j) and augmented contractile profile of cardiospheres without altering the size of the cardiospheres (Fig. 4k-l). Additionally, HB-EGF treatment led to the increase of PDGFRα positive fibroblast areas and collagen deposition in cardiospheres (Fig. 4m-n). Overall, HB-EGF induced a similar phenotype in cardiac fibroblasts and cardiospheres as compared to supernatants of *DNMT3A*-silenced monocytes shown previously in Figure 2, suggesting that it might mediate cardiac fibroblast activation. Indeed, treatment of fibroblasts with supernatants of DNMT3A-silenced monocytes was not only associated with autophosphorylation of the EGFR, but also induced phosphorylation of Akt, the downstream kinase of EGFR signaling, in fibroblasts (Fig. 5a-c). Thus, we can conclude that silencing of DNMT3A in monocytes induces activation of the EGFR signaling pathway.

**Fig. 5:**
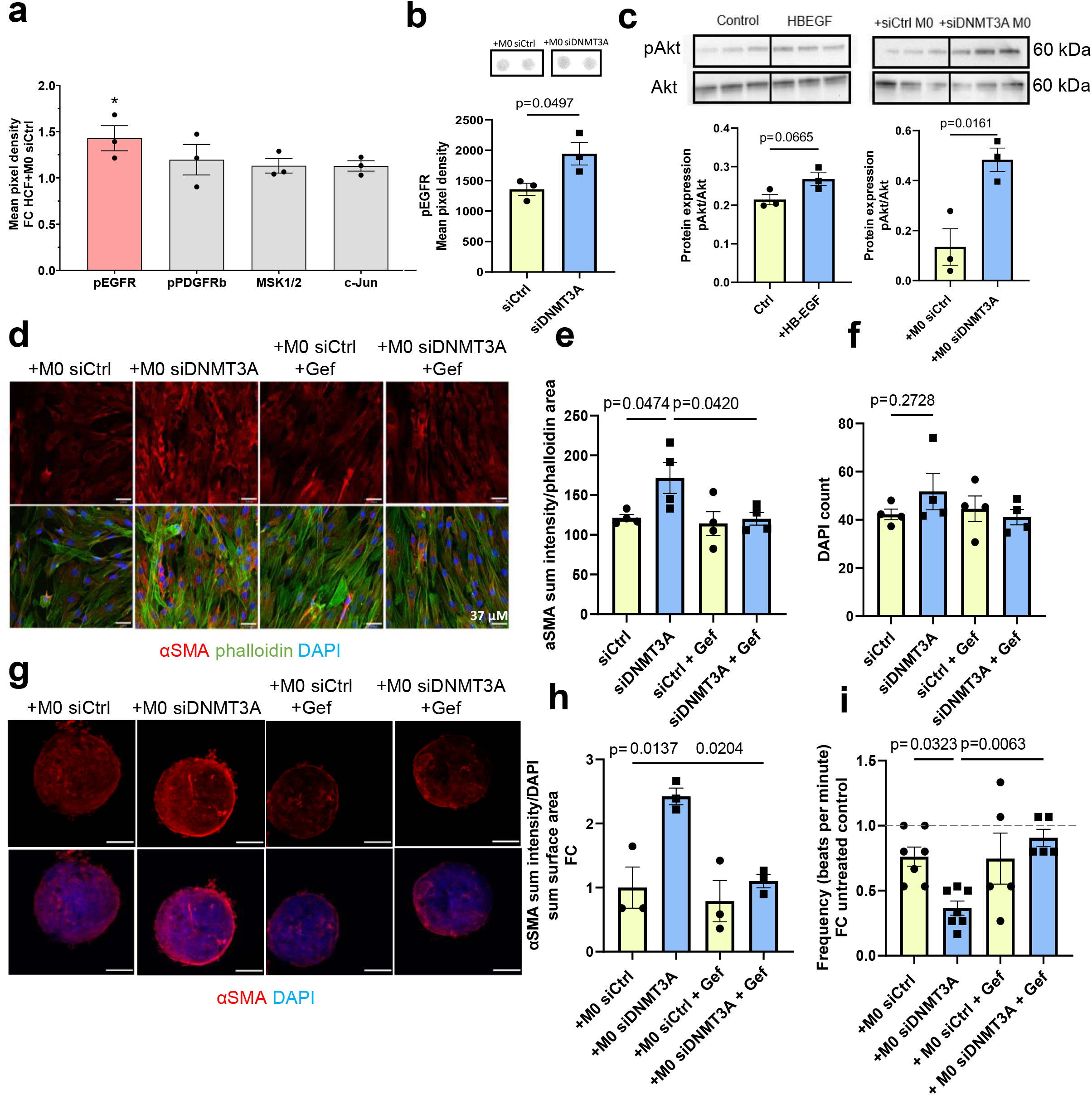
EGF signaling contributes to CHIP monocyte-mediated cardiac fibroblast activation. **a** Alterations in phosphorylated receptors and kinases in HCF treated for 30 min with supernatant from THP1-derived cells with DNMT3A silencing. **b** Quantification of phosphorylated EGFR levels using phospho-kinase array. (n=3) **c** Western Blot for phospho-Akt (pAkt) and total Akt in iHCF stimulated with HB-EGF as a positive control and iHCF stimulated with CHIP supernatants (n=3). **d** Representative immunofluorescence images of *α*SMA protein expression in HCF stimulated with CHIP supernatants and treated with gefitinib. Blue, DAPI; green, phalloidin; red, *α*SMA. **e** Quantification of *α*SMA signal from **d** (n=4). **f** Estimation of cytotoxic effects upon indirect co-culture with HCF and treatment with gefitinib (n=4). **g** Representative immunofluorescence images of *α*SMA protein expression in cardiospheres stimulated with CHIP supernatants and treated with gefitinib. Blue, DAPI; red, *α*SMA. **h** Quantification of *α*SMA signal from **g** (n=4). **i** Estimation of beating frequency cardiospheres stimulated with CHIP supernatants and treated with gefitinib (n=5-7). Data are shown as mean ± SEM. Normal distribution was tested with the Shapiro-Wilk test or the Kolmogorov-Smirnov test. Statistical comparison of four groups was performed using ordinary one-way ANOVA test with posthoc Dunnett’s correction.

Finally, to determine a potential causal involvement of EGF signaling in monocyte-fibroblast crosstalk, we inhibited EGF-receptor signaling in cardiac fibroblasts by the small-molecule EGFR kinase inhibitor gefitinib^35^. While gefitinib did not affect the basal expression of αSMA, it prevented the induction of αSMA by supernatants of DNMT3A-silenced monocytes (Fig. 5d-e), without showing cytotoxic effects in human cardiac fibroblasts (Fig. 5f). Moreover, EGFR inhibition with gefitinib in cardiospheres reduced overexpression of αSMA and abrogated the reduction of cardiac tissue mimetic contractions after treatment with the supernatant of DNMT3A-silenced monocytes (Fig. 5g-i). Taken together, these results demonstrate that the secretome of DNMT3A silenced monocytes induces fibroblasts activation partially through EGFR signaling.

## Discussion

Here, we describe that monocytes derived from patients with heart failure carrying DNMT3A CHIP-driver mutations may contribute to the progression of cardiac fibrosis. Based on the *in silico* predicted increased interaction of monocytes derived from patients harboring DNMT3A CHIP-driver mutations with cardiac fibroblasts, we investigated the paracrine cross-talk and showed that the supernatant of DNMT3A-silenced monocyte activates cardiac fibroblasts in 2D and 3D cultures. Consequently, increased diffuse cardiac fibrosis was confirmed in hematopoietic DNMT3A CHIP-driver mutations carrying mice subjected to myocardial infarction with subsequent heart failure as well as in patients with established heart failure. Together, these data disclose a novel mechanism by which DNMT3A CHIP-driver mutations interfere with heart failure: in addition to the well-known pro-inflammatory activation induced by DNMT3A CHIP-driver mutations, our data point towards a direct interaction of DNMT3A-mutant monocytes with cardiac fibroblasts. As circulating monocytes continuously infiltrate the heart, the activation of cardiac fibroblasts may lead to the instigation and further progression of diffuse cardiac fibrosis in patients with heart failure carrying DNMT3A CHIP-driver mutations.

Mechanistically, we identified HB-EGF as one of the mediators of fibroblast activation. HB-EGF has been implicated in a number of physiological and pathological processes. HB-EGF binds to and activates the EGF receptor (EGFR) via autophosphorylation and induces interstitial fibrosis via the subsequent activation of the Akt/mTor/p70s6k pathway^36,37^. HB-EGF and EGF receptor (EGFR) families are upregulated under pathological conditions such as cardiac hypertrophy or myocardial infarction^38,39^. Shedding and activation or adenoviral overexpression of HB-EGF results in cardiac hypertrophy and increased fibrosis as evidenced by larger αSMA positive areas^37,40^. Consistently, inhibition of HB-EGF or its receptor reduced cardiac pathologies in mouse models^40,41^.

Taken together our data suggesting an augmented fibroblast activation and diffuse fibrosis in DNMT3A CHIP-driver mutation carriers may also have therapeutic implications. Assessment of DNMT3A mutation may identify patients at high risk for negative remodeling and cardiac fibrosis allowing for a targeted treatment with new anti-fibrotic regimens or strategies that specifically interfere with the proposed downstream pathways.

## Materials and methods

### Human single-cell nuclei (sn) and single-cell (sc) RNA sequencing data sets

The following two published datasets were used for analysis: snRNA-seq data from healthy septal cardiac tissue of 14 individuals^17^ and scRNA-seq data from peripheral blood mononuclear cells (PBMCs) of n=5 patients with heart failure (HF) harboring DNA methyltransferase 3A (DNMT3A) clonal hematopoiesis driver mutations (CHIP) and n=4 HF patients without driver mutations (NO-CHIP)^15^.

### Integration and analysis of single-cell nuclei (sn) and single-cell (sc) RNA-sequencing data

For integration and analysis of snRNA- and scRNA-sequencing data, we used Seurat (version 4.0.2). Before performing integration, we retrieved monocytes from transcriptomic dataset of PBMC from CHIP and NO-CHIP carriers^15^ with subset(). To merge snRNA-seq data of cardiac tissue and scRNA-seq datasets of PBMCs, we utilized the merge() function. Expression matrices were normalized and scaled with ‘NormalizeData()’, ‘FindVariableFeatures()’ and ‘ScaleData()’. Using ‘RunPCA()’, we obtained the reduced dimensionality of the merged object. The resulting integration was visualized on a UMAP plot with ‘RunUMAP(). For crosstalk analysis, CellChat (version 1.1.43)^18^ was used. We followed the standard tutorial ‘Comparison analysis of multiple datasets using CellChat’ from the CellChat GitHub repository (https://github.com/sqjin/CellChat).

### Monocytes cell culture and transfection

The human monocytic cell line THP-1 was obtained from the German Collection of Microorganisms and Cell Cultures (#ACC16, DSMZ) and cultured in RPMI 1640 with GlutaMAX medium (#61870036, Thermo Fisher Scientific) supplemented with 10% heat-inactivated fetal bovine serum (FBS) (#10082147, Thermo Fisher Scientific), 10 mM HEPES (#H0887, Sigma Aldrich) and 50 U/mL Penicillin and Streptomycin (#15140122, Thermo Fisher Scientific) at 5% CO2, 37°C.

THP-1 monocytes were differentiated into THP1-derived macrophages by stimulating 5×10^5^ THP-1 monocytes/well in 6-well plate with 100 ng/mL phorbol 12-myristate 13-acetate (PMA) (#P8139, Sigma Aldrich) for 48 h, followed by 24 h of culture in media without PMA. *DNMT3A* (HSS176225) was silenced with 50nM Stealth RNAi siRNAs (#1299001, Invitrogen) by using siTran2.0 transfection reagent (#TT320001, Origene) and 50 nM scrambled siRNA with similar median GC content was used as siRNA control (#12935300, Invitrogen). Transfection was carried out according to the manufacturer’s instructions. The media was changed 24 h post transfection. Supernatants from transfected monocytes were collected 48 h after transfection and used for stimulation.

### Human cardiac fibroblasts cell culture and treatment

Primary human cardiac fibroblasts (HCF) (#C-12375, Promocell) or immortalized human cardiac fibroblasts (iHCF) (#P10453-IM, Innoprot) were used. HCF were and cultured in Fibroblast Growth Medium 3 (#C-23130, Promocell) according to the manufacturer’s protocol. HCF were seeded at a density of 5000 cells/cm^2^ in 6-well, 12-well plates and 8-well μ-Slides (#80826, IBIDI) coated with human fibronectin (#F0895, Sigma Aldrich, 0.1% solution; 1:1000 in water) for indirect co-culture, RNA extraction and qRT-PCR analysis. Immortalized iHCF were cultured in Dulbecco modified Eagles medium (DMEM) (#11965092, Thermo Fisher Scientific) supplemented with 10 % FBS (#16000044, Thermo Fisher Scientific) and 50 U/mL Penicillin and Streptomycin according to the manufacturer’s protocol. HCF were seeded at a density of 17 500 cells/cm^2^ in fibronectin-coated (#F0895, Sigma Aldrich, 0.1% solution; 1:1000 in water) 12-well plates and 8-well μ-Slides for indirect co-culture, RNA extraction and qRT-PCR analysis.

For stimulation with recombinant human HB-EGF (#100-47, PeproTech), fibroblasts were seeded in the respective densities one day prior to the stimulation. One day after stimulation, medium was replaced with fresh medium containing 100 ng/ml HB-EGF for 48 h.

### Human endothelial cell culture

Human umbilical vein endothelial cells (HUVECs, passage 2-3) were purchased from PromoCell (#C-12203 / C-12253) and cultured in endothelial cell basal medium (EBM, #CC-3121, Lonza) supplemented with EGM-SingleQuots (CC-4133, Lonza) and 10% FBS (FCS, #10270-106, Gibco) at 37 °C and 5% CO2. Cells were trypsinized with 0.05% trypsin (#25300062; Thermo Fisher Scientific) for 3 min at 37 °C.

### Indirect co-culture of THP-1 cells and human cardiac fibroblasts

For indirect co-culture experiments, fibroblasts were incubated with the supernatant of transfected THP-1 macrophages. The supernatants were collected 48 h after transfection and were stored at −80°C. THP-1-derived macrophages supernatant was mixed with the respective fibroblast cell medium at a ratio 1:1 (co-culture medium). Fibroblasts were incubated with the mixed medium for 48 hours and then were used for RNA isolation, immunofluorescent staining, and functional assays.

For the EGFR signaling inhibitor experiments, cells were treated with either 5μM gefitinib (#SML1657, Sigma-Aldrich) or water as control. After 2 h, the medium was replaced by co-culture medium and incubated for 48 h with or without addition of the inhibitor to the co-culture medium.

### Western Blot

For Western Blot analysis, protein from transfected THP-1-derived macrophages and human cardiac fibroblasts treated with macrophage supernatant or with recombinant human HB-EGF (100 ng/ml, 30min, #100-47, PeproTech) was isolated. Briefly, cells were washed with ice-cold DPBS (#14190144, Thermo Fischer Scientific) and lysed in RIPA buffer (#R0278, Sigma-Aldrich) supplemented with protease inhibitor cocktail (#P8340, Sigma-Aldrich, 1:50) and phosphatase inhibitor (#P5726, Sigma-Aldrich, 1:100) for 30 min on ice.

Lysates were centrifuged at 12,000 g for 10 min at 4 °C. Protein concentrations were determined using the DC Protein Assay Kit II (#5000112, Bio-Rad). Proteins (20μg) were reduced with Laemmli SDS (6×, #J61337, Alfa Aesar) and were separated on a Mini-PROTEAN TGX gel (4561094, Bio-Rad) (115 V, 50 min) and transferred onto a nitrocellulose membrane using a semi-dry blot (25V, 0,22A, 50 min, Trans-Blot SD Semi-Dry Transfer Cell, #1703940, Biorad) according to the manufacturer’s protocol. Membranes were blocked for 1 h in 5 % Blotto nonfat milk (#sc-2325, Santa Cruz), dissolved in Tris-buffered saline supplemented with Tween-20 (1x TBST, #sc-281695, Santa-Cruz). Primary antibodies were incubated overnight at 4 °C in 5% blocking solution (1:1000 rabbit anti-DNMT3A (E9P2F), #49768, Cell Signaling; 1:1000 rabbit anti-Akt (pan) (C67E7), #4691, Cell Signaling; 1:1000 rabbit anti-phospho-Akt (Ser473) (193H12), #4058, Cell Signaling; 1:1000 rabbit anti-HBEGF (E5L5T), #27450, Cell Signaling; 1:1000 rabbit anti-α-tubulin Antibody, #2144, Cell Signaling). On the next day, membranes were washed with 1x TBST prior to incubation with secondary antibodies. Secondary antibodies were diluted in 5% blocking solution and membranes were incubated for 1 h at room temperature (1:1000 donkey anti-rabbit IgG, HRP-linked Antibody, #7074S, Cell Signaling). Proteins were detected based on HRP substrate-based enhanced chemiluminescence (#WBKLS0500, Millipore), visualized using ChemiDoc Touch Imaging System (BioRad) and quantified using ImageLab (version 5.0).

### Ribonucleic acid analysis

For RNA isolation, cells were lysed with 350 μL RLT Plus buffer and RNA was isolated using the RNeasy Plus Mini Kit according to the manufacturer’s protocol (#74034, Qiagen). RNA content and purity were assessed by spectrophotometry (NanoDrop Technologies). mRNA expression was quantified by by qRT-PCR using 100-1000 ng of total RNA, which was reversed transcribed by MuLV reverse transcriptase (#28025013, Thermo Fisher Scientific) and random hexamer primers (#SO142, Thermo Fisher Scientific). cDNA synthesis was performed according to the protocol: (Primer annealing) 65°C × 5 min, (Primer elongation) 25°C × 10 min, (Reverse transcription) 37°C × 50 min, (Enzyme denaturation) 70°C × 15 min. Expression levels of mRNA were detected by using Fast SYBR Green (#4385612, Applied Biosystems) and an Applied Biosystems Viia7 machine. Cycling conditions on the machine were as follows: (Hold) 95°C × 30 sec, (PCR) 95°C × 1 sec, 60°C × 20 sec repeated for 40 cycles, (Melt curve) 95°C × 15 sec, 60°C × 60 sec, 95°C × 15 sec. Relative gene expression was calculated with the QuantStudio Real-Time PCR software (version 1.3) using 2-^ΔCt^ (^ΔCt^= Ct target gene – Ct housekeeping gene). RPLP0 was used as housekeeping gene.

Human primer sequences are listed below:

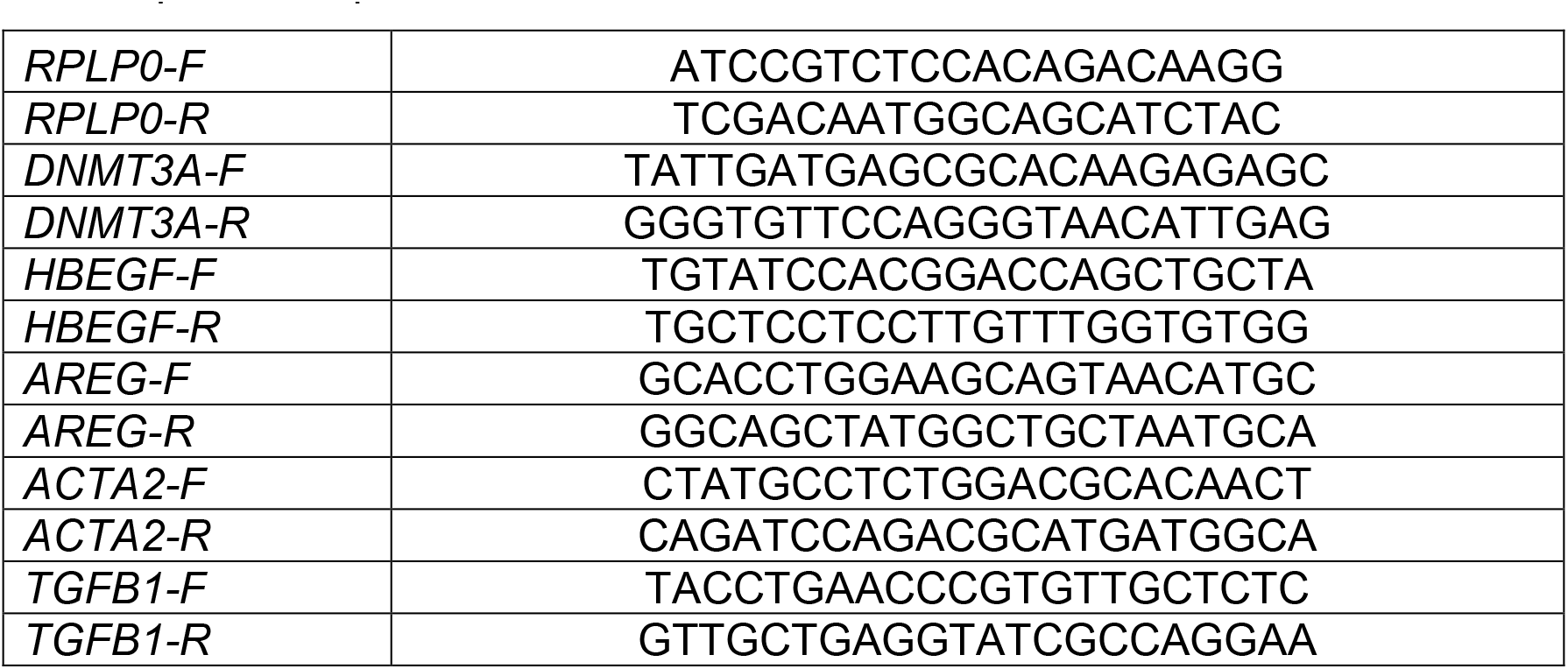

### Immunofluorescence staining for cell culture

μ-Slide with 8 wells (#80826, IBIDI) were pre-coated with human fibronectin (#F0895, Sigma-Aldrich, 0.1% solution) diluted in water 1:1000 for 1 hour at 37°C. Human cardiac fibroblasts were seeded in slides in the respective medium, using HCF with 5·10^3^ cells per well for αSMA staining, 2·10^3^ for phospho-histone H3 staining, and iHCF with 20·10^3^ cells per well. For 48 h after respective treatment, cells were washed two times with DBPS and fixed with 4% PFA (#28908, Thermo Fisher Scientific) for 10 minutes at room temperature. After two washing steps, each for 5 minutes, with DPBS, cells were permeabilized with 0.1 % Triton X-100 (#T8787, Thermo Fisher Scientific) in DPBS for 15 minutes. Cells were blocked with 5% donkey serum (#ab7475, abcam) in DPBS for 60 minutes at RT. Primary antibodies were incubated in the same blocking solution overnight at 4 °C. Cells were stained with phalloidin (1:100, #O7466, Thermo Fisher Scientific), mouse anti-α-smooth muscle actin (1:200, #C6168, Sigma-Aldrich), rabbit anti-phospho-histone H3 (Ser10; 1:200, #06-570, Sigma-Aldrich). Cells were washed 4 times with DPBS, each 5 minutes, and secondary antibodies in DPBS were incubated in for 1 hour at RT. Following dyes and secondary antibodies were used: DAPI (1:1000, #D9542, Merck), donkey anti-rabbit-555 (1:200, #A-31573, Thermo Fisher Scientific). Cells were mounted with Fluoromount-G (#00-4958-0, Invitrogen). Images were taken using Leica STELLARIS confocal microscope. A dry objective was used with a magnification of 20x or objective with oil immersion of 63x. Z-Stack images were acquired with a resolution of 1024×1024 pixels, a speed of 200 frames per second, and a Z-Stack size of approximately 1 μm. Quantification was done using Volocity software version 6.5 (Quorum Technologies).

### Collagen Gel Contraction Assay

Untreated iHCF and macrophage supernatant-treated iHCF (1 × 10 ^5^ per well) were mixed with rat tail collagen type I (#354236, Corning) matrices on a 24-well plate. After 20 min of polymerisation, gels were manually released from the edges of the well and were floating in culture medium only or culture medium mixed with respective supernatants. Gel surface area was measured at 0 h and 24 h with NIS Elements (Nikon Eclipse TS100) and quantified with ImageJ Fiji 2.3.1. Gel contraction was calculated as 1/(gel surface area at 24 h/gel surface area at 0 h) and normalized to the contraction of untreated iHCF.

### Migration Assay

For analysis of cell migration, 2-well culture-inserts for self-insertion (#80209, IBIDI) were put into μ-Slide with 8 wells (#80826, IBIDI) and 7 × 10^4^ iHCF were seeded per division in the inserts. Cells were directly stimulated with mixed medium for 48 hours. Images were taken at 0 h and 8 h after removing the culture-insert with a Nikon Eclipse TS100 microscope. The cell free area was measured using ImageJ Fiji 2.3.1. Subsequently the relative change of the migrative capability was calculated as 1:(cell-free area at 24 hours: cell-free area at 0 hours).

### Cell isolation for cardiac tissue mimetics (cardiospheres)

Neonatal rat cells were isolated from Sprague Dawley P1 and P2 rat pups, as described previously^42^. Mated female Sprague Dawley rats were obtained from Charles River (Sulzfeld, Germany) and Janvier (Le Genest Saint-Isle, France). Rat pups were sacrificed by cervical dislocation and hearts were transferred into Hank’s buffered saline solution (-Ca^2+^/-Mg^2+^; #14025-050, Gibco) containing 0.2% 2,3-Butanedione monoxime (BDM; Sigma-Aldrich; #B0753-25G). Hearts were cut in small pieces and dissociated at 37°C with the commercially available enzyme mix (Neonatal Heart Dissociation Kit, mouse and rat, Miltenyi Biotec GmbH; # 130-098-373) followed by tissue homogenization with the genteMACS^™^ Dissociator (Miltenyi Biotec GmbH; #130-093-235; program m_neoheart_01_01) in a C-tube (#130-093-237, Miltenyi Biotec GmbH). After centrifugation (80 ×g, 5 min), cells were resuspended in plating medium (DMEM high glucose #1-26P02-K, M199 EBS #1-21F01-I; both from BioConcept, 10% horse serum (#16050130; Thermo Fisher Scientific), 5% fetal calf serum (#10270-106; Gibco), 2 % l-glutamine (#25030149; Thermo Fisher Scientific) and penicillin/streptomycin (#11074440001; Roche) and incubated for 1 h and 40 min in 6 cm cell culture dishes (Greiner Bio-One GmbH) at 37°C and 5% CO2. Within the incubation time, the fibroblasts attach to the uncoated culture dish and can be detached by using 0.05 % trypsin (#25300062; Thermo Fisher Scientific) for 3 min at 37 °C. The supernatant contains the cardiomyocytes and was collected. Cells were counted with the Neubauer-Chamber (Carl Roth, #T729.1).

### Cardiac tissue mimetic formation

Isolated rat cardiomyocytes (CM), rat fibroblasts (FB) and HUVECs were used to form cardiac tissue mimetics (CTM) as previously described^26^. Briefly, 32,000 CMs and 6,400 FBs were cultured in plating medium as hanging drops at 37°C and 5% CO2 for 4 days. The formed cellular spheroids were collected and cultivated in Ultra-Low Adhesion U-Bottom plates (#7007, Costar) in 50 μL plating medium per well. After 4h of incubation, 10 000 HUVECs in 50 μL fully supplemented EBM (Lonza) were added to each well. After 24h, the medium was changed to maintenance medium (DMEM high glucose #1-26P02-K, M199 EBS #1-21F01-I; both from BioConcept, 1% horse serum (#16050130; Thermo Fisher Scientific), 2% l-glutamine (#25030149; Thermo Fisher Scientific) and penicillin/streptomycin (#11074440001; Roche) and fully supplemented EBM (50%:50%). Mature CTMs were treated with 200 μM phenylephrine (short PE; Sigma-Aldrich; P6126-5G), every second day for 14 days and then treated PE+ medium and supernatant from macrophages (50%:50%) or PE+ 100 ng/ml recombinant human HB-EGF for 5 days with 3 stimulations. For the inhibitor experiments with gefitinib (#SML1657, Sigma-Aldrich), cardiospheres were treated with PE containing medium and supernatant from macrophages (50%:50%), supplemented with 5 μM gefitinib.

PDGFRα positive areas, αSMA positive areas and vascularisation was determined using the Leica STELLARIS confocal microscope and was quantified with the Volocity software version 6.5 (Quorum Technologies).

### Contractility measurement

Spontaneous cardiospheres contraction were determined by counting the number of beats per minute (bpm) of cardiospheres using a computer-assisted microscope and Axiovision 4.5 (Zeiss).

### Area measurement

Pictures of cardiospheres were taken using a computer-assisted microscope and Axiovision 4.5 (Zeiss). Area of cardiospheres was measured using ImageJ Fiji 2.3.1 (NIH).

### Cardiac tissue mimetics immunofluorescence staining

Whole CTMs were used for fluorescence immunohistochemistry. CTMs were collected and fixed with 4% ROTI Histofix (#P087.1, Carl-Roth GmbH & Co. KG) for 1h at room temperature. After fixation, CTMs were permeabilized with 0.2% Triton X-100 (#T8787, Thermo Fisher Scientific) in DPBS for 1 h at room temperature followed by incubation in blocking solution (2% donkey serum (#ab7475, abcam), 3% BSA (#A7030, Sigma-Aldrich) in 0.2% Triton X-100 in DBPS) for 30 min at room temperature.

Primary antibodies and dyes (mouse anti-α-smooth muscle actin (1:40, #C6168, Sigma-Aldrich), goat anti-PDGFRα (1:20, #AF1062, R&D Systems), biotinylated Ulex Europaeus Agglutinin I (1:100, #B-1065-2, Vector Laboratories) were diluted in blocking solution and were incubated overnight at 4°C. CTMs were washed on the next day with 0.2% Triton X-100 in DBPS three times for 20 min. Secondary antibodies and dyes (streptavidin 488 (#S11223, Thermo Fisher Scientific), donkey anti-goat (1:200 #A-21447, Thermo Fisher Scientific), DAPI (1:1000, #D9542, Merck) were diluted in the blocking solution and incubated for 4h at room temperature in the dark. After washing CTMs again three times with 2% Triton X-100 in DBPS, CTMs were mounted with Fluoromount-G (#00-4958-0, Invitrogen) and analyzed using the Leica STELLARIS confocal microscope with quantification performed in Volocity software version 6.5 (Quorum Technologies).

### Generation of mouse DNMT3A CHIP model

Mx-Cre^+^/Dnmt3a (Ly5.2) and Mx-Cre^-^Dnmt3a (Ly5.2) have been previously described^28^. B6.SJL (Ly5.1) mice were obtained from The Jackson Laboratory. Animals were housed under standard laboratory conditions. Age-matched Mx-Cre^+^/Dnmt3a and Mx-Cre^-^/Dnmt3a donor mice (Ly5.2) were treated with pI:pC (400μg/mouse intraperitoneally [i.p.]) for 3 nonconsecutive days (Amersham). Donor bone marrow cells (2×10^6^ cells) were intravenously injected into lethally irradiated (7.5Gy total body irradiation) B6.SJL recipients (Ly5.1). Recipient mice were maintained on antibiotic-containing drinking water (ciprofloxacin 50 mg/kg) 5 days pre-lethally irradiation and 2 weeks post irradiation and transplantation. Peripheral blood engraftment was assessed by flow cytometry 6 weeks after reconstitution.

### Left anterior descending artery (*LAD*) ligation in mice

Six weeks after reconstitution of the bone marrow, myocardial infarction was induced by permanent ligation of the left anterior descending artery in female mice as described previously^27^. In brief, anesthesia was induced with isoflurane (4%/800 ml O_2_/min) and maintained by endotracheal ventilation (2–3%/800 ml O_2_/min). Thoracotomy was performed in the fourth left intercostal space. The left ventricle was exposed, and the left coronary artery was permanently occluded. Chest and skin were closed, and anesthesia was terminated. Animals were extubated when breathing was restored. Initial myocardial injury was evaluated by measuring cardiac troponin-T levels in plasma 24 h after induction of myocardial infarction.

### Collagen deposition analysis

To analyze fibrotic scar area and diffuse fibrosis, we checked collagen deposition in the heart by performing Sirius Red staining on paraffin sections. After deparaffinization and rehydration, the slides were stained with 0.1 % Picro Sirius Red solution prepared using Sirius Red F3BA (#1A280, Waldeck GmbH) in an aqueous solution of Picric Acid (#6744, Sigma Aldrich). Slides were mounted with Aquatex mounting medium (#HC440258, Millipore) and imaged using a Nikon Eclipse Ci microscope with a 2X (fibrotic scar) and 40X (diffuse fibrosis) objectives. Positive area was measured using ImageJ (NIH, version 1.53a) and normalized either to the heart area (scar analysis) or to the total tissue area (diffuse fibrosis).

### Cardiac magnetic resonance (CMR) imaging and post-processing

All participants underwent a standardized CMR imaging protocol at the Centre for Cardiovascular Imaging, University Frankfurt, on a 3T scanner equipped with advanced cardiac software and a multi-channel coil (Skyra and Prisma, Siemens Healthineers, software version VE11) as previously described^30,31^. The study was approved by an institutional review committee of the University Hospital of the Johann Wolfgang Goethe University in compliance with internal standards of the German government, and procedures followed were in accordance with institutional guidelines (application 347/18) and the Declaration of Helsinki. Acquisition of cardiac function, volumes, mass, myocardial mapping and scar imaging was performed. Imaging parameters and scanning and shimming procedures for all sequences were standardized and mandatorily performed by all operators in all scans. Comparability and reproducibility of measurements were determined at each location. Slice thickness in all acquisitions was set uniformly at 8 mm. Cine imaging was performed using a balanced steady-state free precession sequence in combination with parallel imaging (GRAPPA) and retrospective gating during expiratory breath-hold (TE/TR/flip-angle: 1.7 ms/3.4 ms/30°; spatial resolution, 1.8 × 1.8 ×8 mm), as a short-axis (SAX) stack for assessment of cardiac volumes and function or single-slice long-axis views (two-chamber, three-chamber and four-chamber views). Cardiac volumes, function and mass were measured in line with standardized post-processing recommendations^30,31^. Cine images were employed for derivation of GLS analyses using Medis Suite MR version 2.1 (Medis Medical Imaging Systems). Myocardial mapping was acquired in a single midventricular slice and measured conservatively in the septal myocardium^8^.

### ELISA

Peripheral blood samples were taken from heart failure with reduced ejection fraction (HFrEF) patients at the Department of Cardiology, University Frankfurt, in the frame of the UCT-Project-Nr.: KardioBMB#2022-004. The study was approved by an institutional review committee of the University Hospital of the Johann Wolfgang Goethe University in compliance with internal standards of the German government, and procedures followed were in accordance with institutional guidelines (application 347/18) and the Declaration of Helsinki. Aliquots of serum and whole blood samples were collected and stored at −80°C. Serum was diluted 1:4 prior to the analysis. Diluted serum was then used to measure human HB-EGF with the DuoSet ELISA Kit (DY259B, R&D Systems).

### Next-generation sequencing

Whole blood from patients with heart failure with reduced ejection fraction (HFrEF) was used for DNA isolation and next-generation sequencing as previously described^10^. Briefly, next-generation sequencing was commercially performed by MLLDxGmbH, Munchen, Germany. DNA was isolated with the MagNaPure System (Roche Diagnostics, Mannheim, Germany) from mononuclear cells after lysis of erythrocytes. The patients’ libraries were generated with the Nextera Flex for enrichment kit (Illumina, San Diego,CA, USA) and sequences for DNMT3A enriched with the IDT xGen hybridization capture of DNA libraries protocol and customized probes (IDT, Coralville, IA, USA). The libraries were sequenced on anIllumina NovaSeq 6000 with a mean coverage of 2147x and a minimum coverage of 400x. Reads were mapped to the reference genome (UCSC hg19) using Isaac aligner (v2.10.12) and a small somatic variant calling was performed with Pisces (v5.1.3.60).

### Human Phospho-Kinase Array

The human Proteome Profiler Human Phospho-Kinase Array Kit (#ARY003C, R&D Systems) was used to assess receptor phosphorylation and kinase activation of HCF treated with macrophages supernatants for 30 min. Briefly, HCF were stimulated with mixed medium in a 6-well plate, lysed in 334 μl of a provided lysis buffer and processed according to manufacturer’s protocol.

### Statistical analysis

Data are biological replicates and are represented as mean and error bars indicating standard error of the mean (SEM). Data were statistically assessed for Gaussian distribution using Shapiro-Wilk, Kolmogorov-Smirnov and Anderson-Darling test. Statistical significance for data with a Gaussian distribution was calculated using two-sided, unpaired Student’s t-tests for two-group comparison. For data not following a Gaussian distribution, statistical analysis was performed using Wilcoxon-Mann-Whitney test for two-group comparison. For multiple comparisons, ordinary one-way ANOVA with a post hoc Tukey’s, Holm-Sidak’s or Dunett’s multiple comparison (following Gaussian distribution) or a Kruskal-Wallis test with a post hoc Dunn’s multiple comparison (not following Gaussian distribution) was used. Fisher’s exact test was used in the analysis of contingency tables. All calculations were performed in GraphPad Prism 9.3.0.

## Data availability

The authors declare that the data underlying the findings of this study are available within the paper or are available upon request per.

## Acknowledgements

SD is supported by the Dr. Robert Schwiete Stiftung and the German Research Foundation (DFG; SFB 1366, Project B04). AMZ is supported by the ERC advanced grant CHIP-AVS Zeiher (Project number 101054899), and DZHK Cellular Heterogeneity. SC is supported by the DFG (SFB 1531, Project number 456687919; Project B10). VP, EN, IH are supported by the DZHK.

## Author contributions

M.S., G.L., A.M.Z. and S.D. designed research; M.S., V.P., I.H., B.S., A.D., S.F.G, M.M.R, D.J. performed research; V.P, E.N., J.H. provided samples and CMR data of patients with heart failure; SC and KK provided serum of patients with HF and DNMT3A mutations; I.H., X.L., M.Sc, C.M.T and F.L. provided samples of WT and DNMT3A CHIP mice; M.S., V.P., W.T.A., J.H., D.J., S.C. and S.D. analyzed data; M.S., G.L., A.M.Z. and S.D. wrote the paper.

## Competing interests

The authors declare no competing interests.

## Materials & Correspondence

Correspondence and material requests should be addressed to Prof. Stefanie Dimmeler.

---

**Supplementary Table 1.** Baseline characteristics of the cMRI study cohort. Baseline characteristics

Baseline characteristics
|  | Characteristic | Total cohort (n = 38) | NO-CHIP (n = 32) | CHIP (n = 6) | P value (CHIP versus NO CHIP) |
| --- | --- | --- | --- | --- | --- |
| 1 | Age, mean (SD) (n = 38) | 66.34 (9.49) | 66.25 (9.99) | 66.83 (6.85) | 0.8923 |
| 2 | Male, No. (%) (n = 38) | 30 (78.95) | 26 (81.25) | 4 (66.67) | 0.587 |
| 3 | Weight (kg), mean (SD) (n = 38) | 82.47 (16.17) | 82.81 (16.15) | 80.67 (17.70) | 0.77 |
| 4 | Size (cm), mean (SD) (n = 38) | 174.1 (10.96) | 173.5 (10.93) | 177.5 (11.52) | 0.416 |
| 5 | Hypertension, No. (%) (n = 34) | 32 (94.12) | 27 (96.43) | 5 (83.33) | 0.3262 |
| 6 | Diabetes mellitus, No. (%) (n = 14) | 6 (42.86) | 4 (40.00) | 2 (50.00) | >0.9999 |
| 7 | Smoking, No. (%) (n = 36) | 24 (66.67) | 19 (63.33) | 5 (83.33) | 0.6399 |
| 8 | Valve disease, No. (%) (n = 32) | 6 (18.75) | 5 (18.52) | 1 (20.00) | >0.9999 |
| 9 | Family history of coronary artery disease, No. (%) (n = 14) | 7 (36.84) | 6 (42.86) | 1 (20.00) | 0.6027 |
| 10 | Coronary artery disease, No. (%) (n = 35) | 16 (45.71) | 14 (48.28) | 2 (33.33) | 0.4227 |
| 11 | Myocardial infarction, No. (%) (n = 36) | 7 (19.44) | 5 (16.67) | 2 (33.33) | 0.5732 |
| 12 | Myocarditis, No. (%) (n = 35) | 1 (2.86) | 0 (0) | 1 (16.67) | 0.1714 |
| 13 | NYHA class, mean (SD) (n=33) | 1.818 (0.7687) | 1.926 (0.7808) | 1.333 (0.5164) | 0.1056 |
| 14 | NT-proBNP serum levels (pg/ml), mean (SD) (n = 31) | 8869 (18763) | 7540 (16540) | 14406 (27450) | 0.7206 |
| 15 | High sensitivity C-reactive protein levels (mg/dl), mean (SD) (n = 31) | 38.45 (64.43) | 43.77 (69.15) | 10.8 (10.5) | 0.1466 |
| 16 | High sensitive troponin T levels (pg/ml), mean (SD) (n = 32) | 137.1 (294.3) | 123 (289) | 197.8 (338) | 0.1768 |
| 17 | Hemoglobin (g/dl), mean (SD) (n = 33) | 128.4 (41.65) | 133.4 (37.91) | 105.5 (53.50) | 0.1109 |
| 18 | Hematocrit, mean (SD) (n = 33) | 374.8 (120.2) | 392.6 (109.3) | 295.2 (144.8) | **0.0497** |
| 19 | Thrombocytes (/μl), mean (SD) (n = 32) | 245.2 (63.83) | 235.8 (61.47) | 285.8 (62.65) | 0.1055 |
| 20 | Leukocytes (/μl), mean (SD) (n = 33) | 618.3 (267.9) | 677 (211.9) | 354 (351.5) | **0.0055** |
| 21 | ACE inhibitor /AT1 blocker, No. (%) (n = 32) | 18 (56.25) | 13 (50.00) | 5 (83.33) | 0.1959 |
| 22 | Aldosterone blocker, No. (%) (n = 32) | 18 (56.25) | 14 (53.85) | 4 (66.67) | 0.6722 |
| 23 | Betablocker, No. (%) (n = 32) | 27 (84.38) | 21 (80.77) | 6 (100) | 0.5546 |
| 24 | Statin, No. (%) (n = 32) | 21 (65.63) | 17 (65.38) | 4 (66.67) | >0.9999 |
| 25 | Diuretics, No. (%) (n = 32) | 13 (40.63) | 9 (34.62) | 4 (66.67) | 0.1937 |

**Supplementary Table 1.** cd. Baseline characteristics of the cMRI study cohort. cMRI findings

cMRI findings
|  | Characteristic | Total Cohort (n = 38) | NO-CHIP (n = 32) | CHIP (n = 6) | P value (CHIP versus NO CHIP) |
| --- | --- | --- | --- | --- | --- |
| 1 | LVEF (%), mean (SD) (n = 35) | 34.06 (9.11) | 34.83 (9.22) | 30.33 (8.287) | 0.5561 |
| 2 | LV-EDVi (mL/m2), mean (SD) (n = 33) | 113.6 (40.14) | 111.1 (40.3) | 124.5 (41.15) | 0.4699 |
| 3 | LV mass index (g/m2), mean (SD) (n = 32) | 73.03 (31.53) | 75.23 (33.76) | 63.5 (18.2) | 0.3864 |
| 4 | RVEF (%), mean (SD) (n = 34) | 43.26 (11.14) | 44.18 (11.49) | 39 (8.922) | 0.3085 |
| 5 | Native T1 (ms), mean (SD) (n = 33) | 1146 (96.64) | 1131 (97.46) | 1214 (61.15) | **0.0497** |
| 6 | Native T2 (ms), mean (SD) (n = 33) | 39.42 (4.10) | 39.41 (4.00) | 39.5 (4.93) | 0.7754 |
| 7 | Pericardial effusion, No. (%) (n = 35) | 11 (31.42) | 8 (26.67) | 3 (60.00) | 0.297 |
| 8 | LGE, No (%) (n = 30) | 21 (70.00) | 17 (70.83) | 4 (66.67) | >0.9999 |

**Supplementary Table 2.**
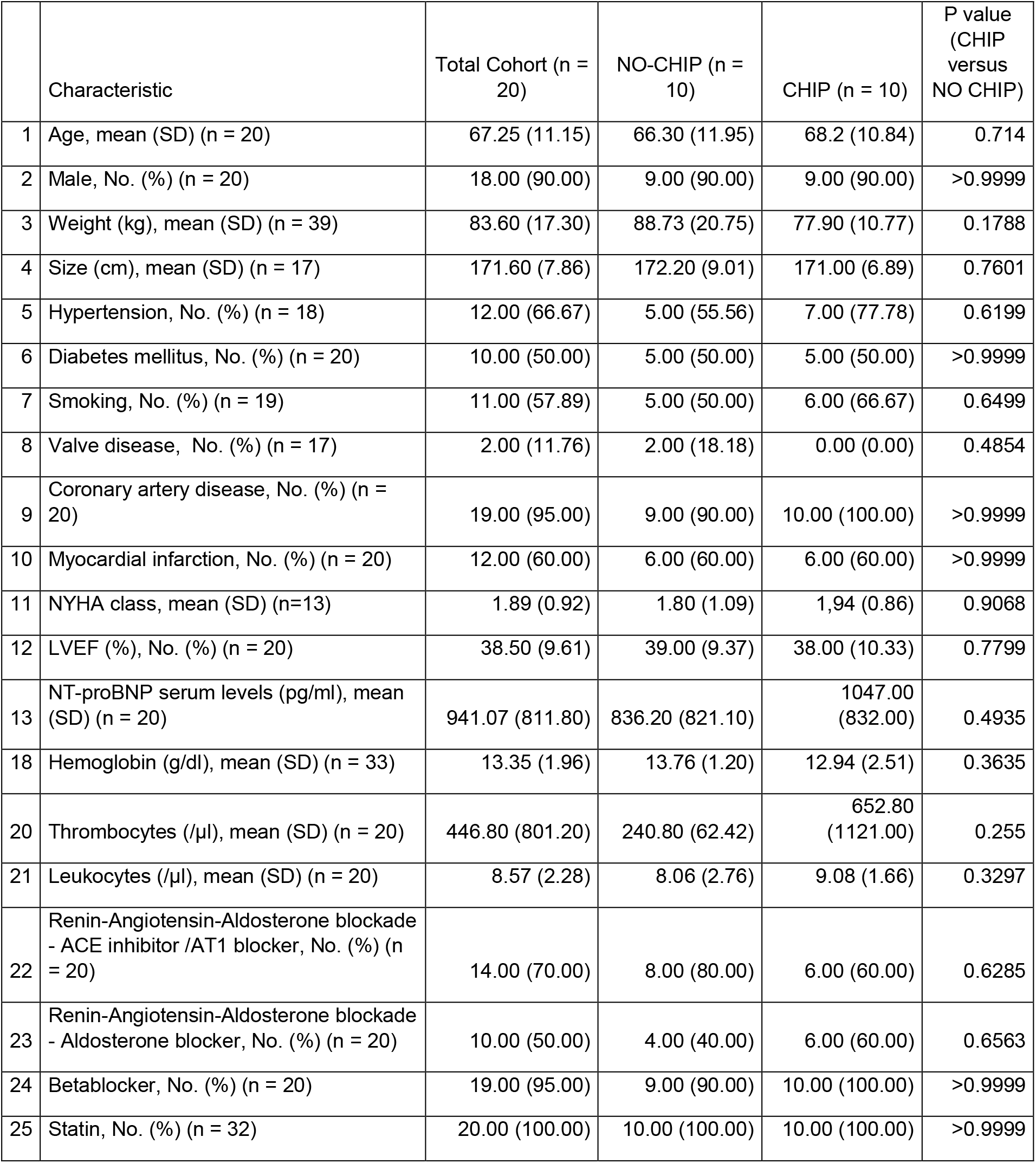
Baseline characteristics of the serum study cohort.

|  | Characteristic | Total Cohort (n = 20) | NO-CHIP (n = 10) | CHIP (n = 10) | P value (CHIP versus NO CHIP) |
| --- | --- | --- | --- | --- | --- |
| 1 | Age, mean (SD) (n = 20) | 67.25 (11.15) | 66.30 (11.95) | 68.2 (10.84) | 0.714 |
| 2 | Male, No. (%) (n = 20) | 18.00 (90.00) | 9.00 (90.00) | 9.00 (90.00) | >0.9999 |
| 3 | Weight (kg), mean (SD) (n = 39) | 83.60 (17.30) | 88.73 (20.75) | 77.90 (10.77) | 0.1788 |
| 4 | Size (cm), mean (SD) (n = 17) | 171.60 (7.86) | 172.20 (9.01) | 171.00 (6.89) | 0.7601 |
| 5 | Hypertension, No. (%) (n = 18) | 12.00 (66.67) | 5.00 (55.56) | 7.00 (77.78) | 0.6199 |
| 6 | Diabetes mellitus, No. (%) (n = 20) | 10.00 (50.00) | 5.00 (50.00) | 5.00 (50.00) | >0.9999 |
| 7 | Smoking, No. (%) (n = 19) | 11.00 (57.89) | 5.00 (50.00) | 6.00 (66.67) | 0.6499 |
| 8 | Valve disease, No. (%) (n = 17) | 2.00 (11.76) | 2.00 (18.18) | 0.00 (0.00) | 0.4854 |
| 9 | Coronary artery disease, No. (%) (n = 20) | 19.00 (95.00) | 9.00 (90.00) | 10.00 (100.00) | >0.9999 |
| 10 | Myocardial infarction, No. (%) (n = 20) | 12.00 (60.00) | 6.00 (60.00) | 6.00 (60.00) | >0.9999 |
| 11 | NYHA class, mean (SD) (n=13) | 1.89 (0.92) | 1.80 (1.09) | 1.94 (0.86) | 0.9068 |
| 12 | LVEF (%), No. (%) (n = 20) | 38.50 (9.61) | 39.00 (9.37) | 38.00 (10.33) | 0.7799 |
| 13 | NT-proBNP serum levels (pg/ml), mean (SD) (n = 20) | 941.07 (811.80) | 836.20 (821.10) | 1047.00 (832.00) | 0.4935 |
| 18 | Hemoglobin (g/dl), mean (SD) (n = 33) | 13.35 (1.96) | 13.76 (1.20) | 12.94 (2.51) | 0.3635 |
| 20 | Thrombocytes (/μl), mean (SD) (n = 20) | 446.80 (801.20) | 240.80 (62.42) | 652.80 (1121.00) | 0.255 |
| 21 | Leukocytes (/μl), mean (SD) (n = 20) | 8.57 (2.28) | 8.06 (2.76) | 9.08 (1.66) | 0.3297 |
| 22 | Renin-Angiotensin-Aldosterone blockade - ACE inhibitor /AT1 blocker, No. (%) (n = 20) | 14.00 (70.00) | 8.00 (80.00) | 6.00 (60.00) | 0.6285 |
| 23 | Renin-Angiotensin-Aldosterone blockade - Aldosterone blocker, No. (%) (n = 20) | 10.00 (50.00) | 4.00 (40.00) | 6.00 (60.00) | 0.6563 |
| 24 | Betablocker, No. (%) (n = 20) | 19.00 (95.00) | 9.00 (90.00) | 10.00 (100.00) | >0.9999 |
| 25 | Statin, No. (%) (n = 32) | 20.00 (100.00) | 10.00 (100.00) | 10.00 (100.00) | >0.9999 |

**Supplementary figure 1.**
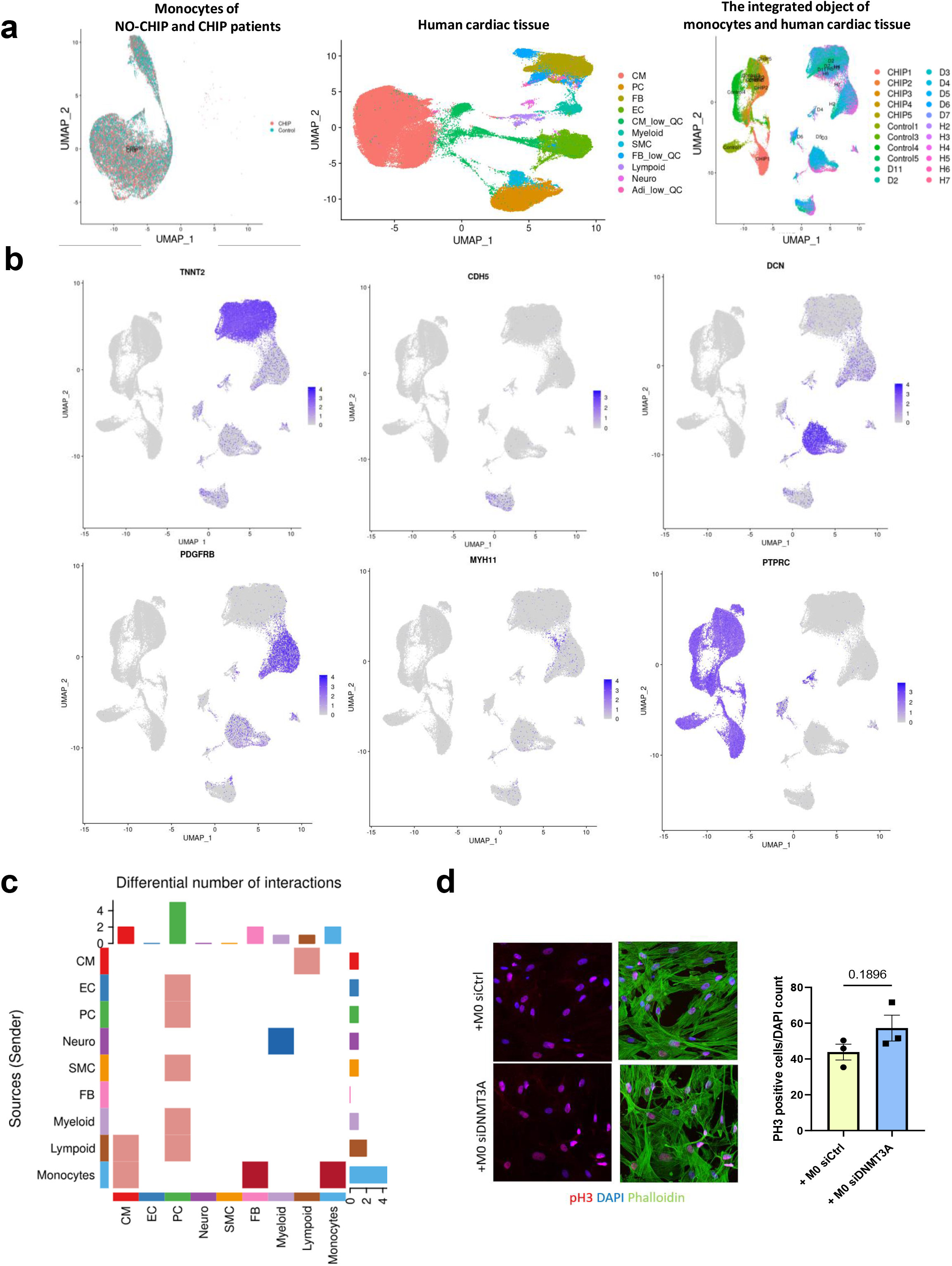
scRNA-seq and scnRNA-seq quality control and monocytes/macrophages-to-fibroblasts interactions. **a** Representative uniform manifold approximation and projection (UMAP) plots before and after integration – scRNA-seq dataset from PBMC obtained from HF patients with DNMT3A CHIP and without CHIP showing samples, conditions, cell types. Left, monocytes dataset from PBMCs of NO-CHIP and CHIP patients before integration, sorted in conditions. Middle, human cardiac tissue dataset before integration. Right, the integrated object of the monocytes and human cardiac tissue, showing individual samples **b** Feature plots for established cell-type specific marker genes: *TNNT2* for CM, *CDH5* for EC, *DCN* for FB, *PDGFRB* for PC, *MYH11* for SMC, *PTPRC* for leukocytes. **c** Representative uniform manifold approximation and projection (UMAP) Result of cellular crosstalk analysis with CellCat depicting differential number of interactions between celltypes. Red, upregulated in CHIP condition. Blue, downregulation of CHIP interactions. **d** Left, representative immunofluorescence images of phospho-histone H3 protein expression in supernatant-stimulated HCF. Blue, DAPI; green, phalloidin; red, pH3. Right, quantification of a pH3 positive nuclei (n=3). Data are shown as mean ± SEM. Normal distribution was tested with the Shapiro-Wilk test or the Kolmogorov-Smirnov test. Statistical comparison of two normally distributed groups was performed using unpaired, two-sided Student’s t-test.

**Supplementary Figure 2.**
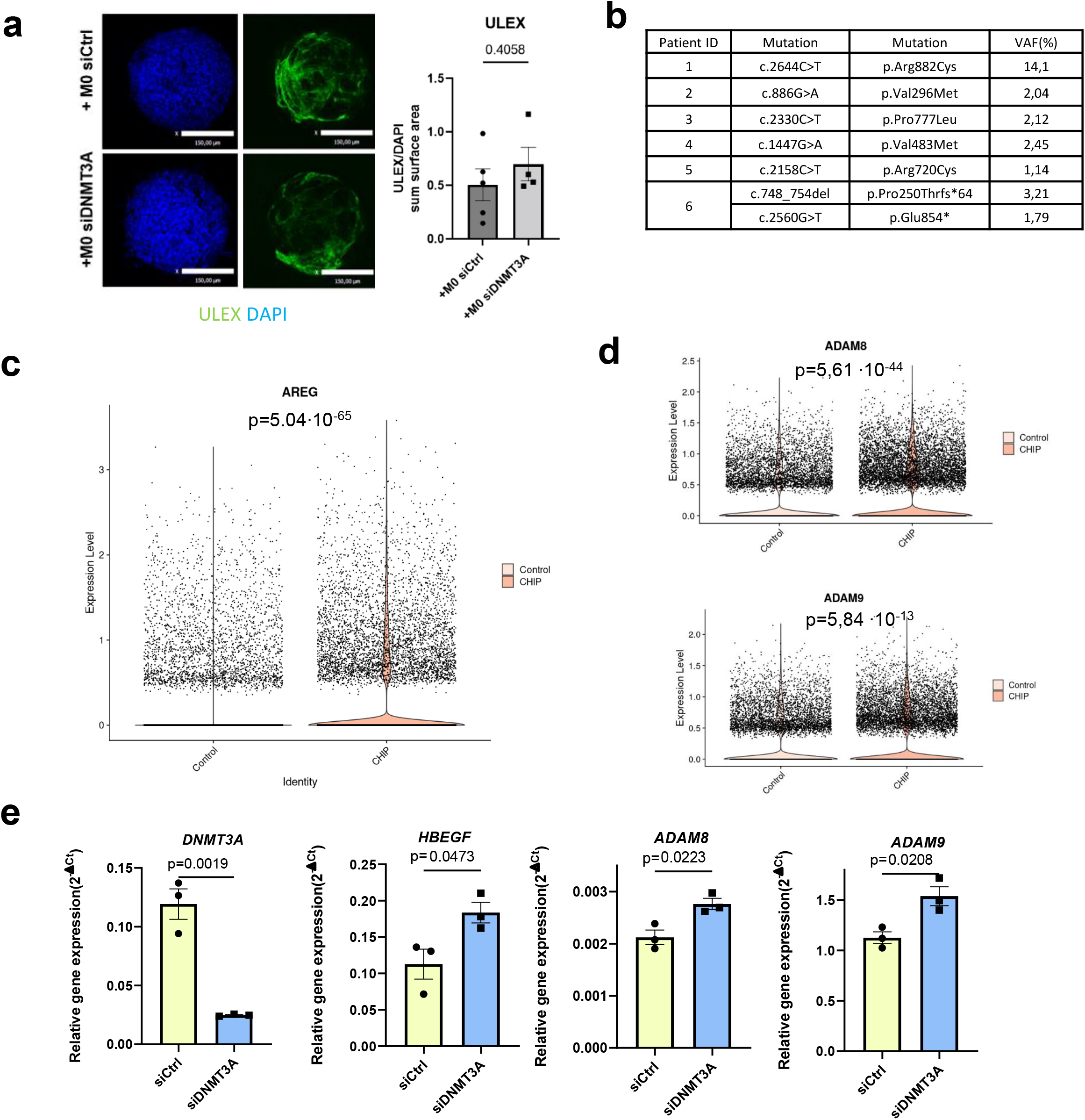
DNMT3A CHIP monocytes/macrophages interact with fibroblasts through EGFR pathway. **a** Left, representative immunofluorescence images of vessel density in stimulated cardiospheres. Blue, DAPI; green, ULEX. Right, quantification of ULEX positive area (n=4-5). **b** DNMT3A CHIP mutation profile for 6 HF patients in the cMRI cohort; VAF - variant allele frequency. **c** Violin plot for *AREG* gene expression from scRNA-seq data of monocytes from No-CHIP and CHIP HF patients. **d** Violin plots showing *ADAM8* and *ADAM9* gene expression from scRNA-seq data of monocytes from No-CHIP and CHIP HF patients. **e** Transcript expression of *DNMT3A*, *HBEGF*, *ADAM8* and *ADAM9* in DNMT3A-silenced THP-1 monocytes (n=3). Normal distribution was tested with the Shapiro-Wilk test or the Kolmogorov-Smirnov test. Statistical comparison of two normaly distributed groups was performed using unpaired, two-sided Student’s t-test

## Notes

### Competing Interest Statement

The authors have declared no competing interest.

